# Strategy-dependent causal relationships between large-scale cortical activity and perceptual decisions

**DOI:** 10.64898/2026.10.31.763327

**Authors:** Zeinab Mohammadi, Jonathan W. Pillow, Lucas Pinto, Joshua I. Glaser

**Affiliations:** Department of Neurology, Northwestern University, Chicago IL 60611; Department of Computer Science, Northwestern University, Evanston IL 60208; Princeton Neuroscience Institute, Princeton University, Princeton NJ 08540; Department of Neurobiology, The University of Chicago, Chicago IL 60637; National Institute for Theory and Mathematics in Biology, Chicago, IL, 60611, USA

**Author notes:** Shared senior authorship (Lucas Pinto, Joshua I. Glaser).

## Abstract

Animals can adopt different internally generated strategies to solve decision-making tasks, but whether cortical regions distinctly contribute to decisions under different strategies remains unknown. To address this problem, we analyzed three datasets comprising either cortical widefield imaging, or optogenetic inactivation of dozens of cortical locations in 54 mice, all performing the same sensory-evidence-accumulation task. Using Generalized Linear Model-Hidden Markov Models (GLM-HMMs), we found that the mice adopted one of three underlying strategies: an engaged, stimulus-driven state, and two disengaged states characterized by large side biases. The specific effects of inactivation of cortical regions depended on behavioral strategy: during engaged states, inactivation tended to increase side bias, while the opposite was true in biased states. For engaged states, these effects were most prominent when frontal regions were inactivated. Moreover, neural decoding analyses from cortex-wide activity revealed that frontal regions preferentially encode the animals’ choices during engaged states relative to biased states. Our results suggest that distinct patterns of large-scale cortical dynamics underlie decisions under different task strategies and identify the frontal cortex as an important substrate of engaged-strategy control.

## 1 Introduction

When performing decision-making tasks, rodents can switch between different strategies, even within the same behavioral session. For example, during perceptual decisions, they may adopt the optimal stimulus-driven decision policy and/or degenerate strategies like choice perseveration or relying on reward and stimulus history [1–15], or even change between different embodiment strategies [14, 16, 17]. However, we understand little about the neural mechanisms that implement and switch between these internally generated strategies. This is for two main reasons. First, most previous studies of decision making have implicitly assumed the existence of a fixed strategy. Second, work on the neural mechanisms of decision-making strategies has primarily happened at the behavioral level or employed neural recordings [11–15], meaning that we lack a clear causal picture. On the other hand, optogenetic perturbation studies have shown that the dorsomedial striatum and a few areas of the dorsal cortex (premotor, posterior parietal, visual area P) have strategy-dependent contributions to behavior [9, 10, 16], suggesting that different strategies causally rely on distinct large-scale activity patterns. This remains untested because previous studies have focused on single or few brain regions.

To address this, here we developed a modeling approach to integrate published datasets consisting of a large optogenetic-silencing and neural-activity survey of regions across the dorsal cortex from dozens of mice performing a visual evidence-accumulation task in virtual reality [18, 19]. We used generalized linear-hidden Markov models (GLM-HMMs) to infer a small number of behavioral strategies—discrete latent states that transition following a Markov process, with each state defined by its own mapping from task covariates to choice [11–13, 20–27]—asking how neural activity and its causal relationship are strategy dependent. While the effects of inactivation during the task are widespread, as we have previously shown, we here discovered that these effects are qualitatively different depending on whether the mice were using stimulus-driven engaged strategies or sub-optimal, side-biased ones. In particular, neural-coding properties and silencing effects were most prominent for frontal regions during the engaged strategy. Together, our findings provide a novel characterization of how large-scale cortical dynamics are reconfigured to have distinct causal links to choice depending on internally generated decision-making strategies.

## 2 Results

To understand cortex-wide links to behavioral strategies, we leveraged multiple datasets in which mice performed the same virtual-reality decision-making task—the “accumulating-towers” task (Fig. 1a). Briefly, mice navigating through a virtual T-maze encountered pulsed visual cues (“towers”) on either side during the first 2 m, with variable counts and positions across trials. After a cue-free 1-m delay, they were rewarded for turning into the arm on the same side with the highest number of cues [6]. The datasets encompassed optogenetic perturbations at different temporal scales, along with widefield calcium imaging. More precisely, the three datasets included 1) whole-trial optogenetic inactivations across 29 dorsal cortical locations [18] (*n* = 20 mice, 178,430 trials); 2) sub-trial inactivations across 9 cortical locations [19] (*n* = 28 mice, 239,351 trials); and 3) widefield calcium imaging from across the dorsal cortex [18] (*n* = 6 mice, 5,321 trials) (Fig. 1b). Because the task was identical across the datasets, we were able to jointly model the underlying behavioral strategies in each of them, while distinctly determining how perturbations and neural activity related to those strategies.

**Fig. 1:**
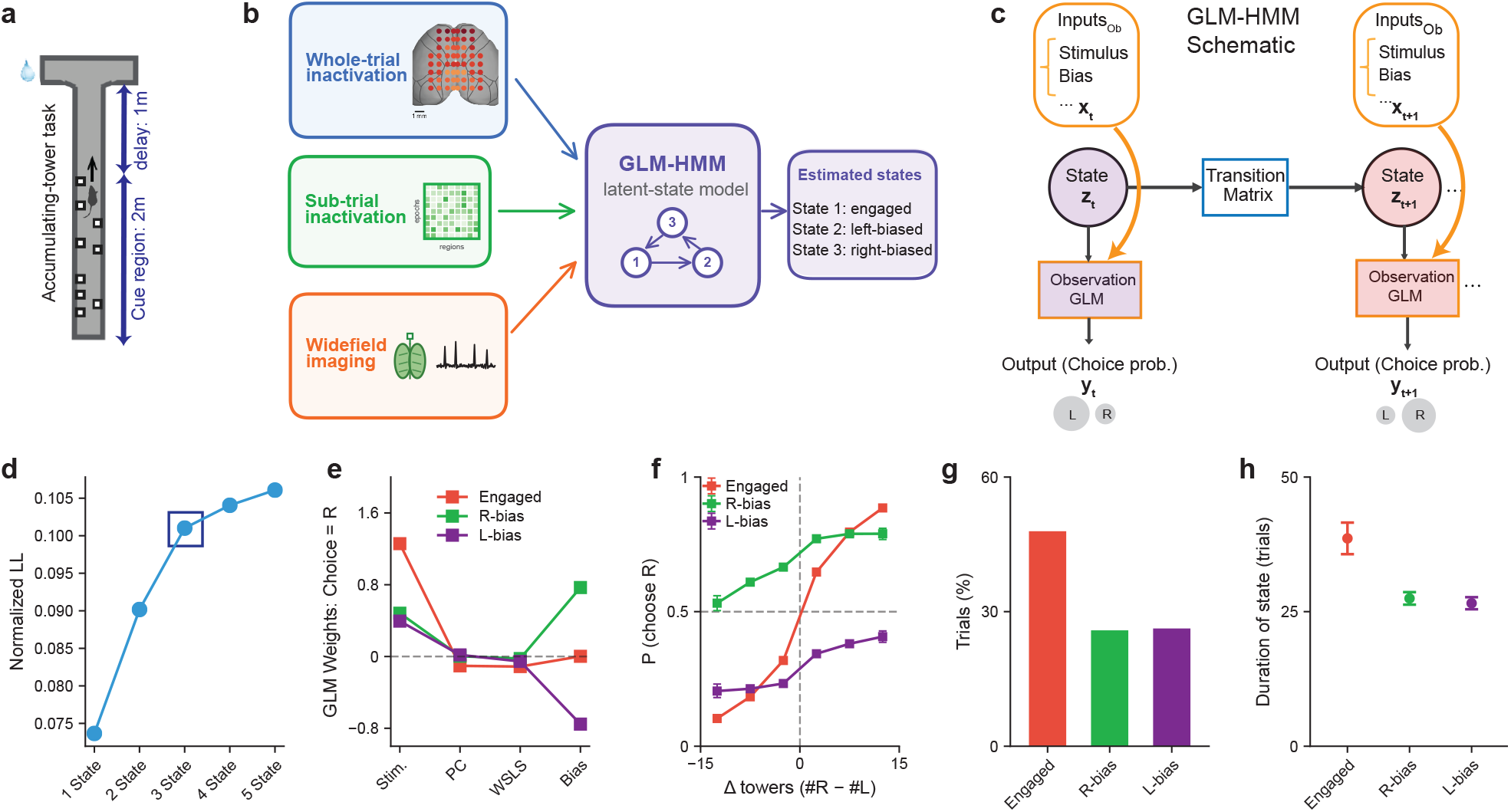
A GLM-HMM fit to behavioral data across datasets reveals three persistent decision-making states. **(a)** In the virtual-reality accumulating-towers task [6], mice viewed visual cues along the first 2 m of the maze, followed by a 1-m cue-free delay, before choosing a turn at the T-junction. **(b)** We fit a GLM-HMM to pooled behavioral decision-making (choice) data from three datasets, yielding three states: engaged, left-biased, and right-biased. **(c)** GLM-HMM framework: state-specific GLMs use covariates such as sensory evidence and previous choice to predict the choice on each trial, with a unique GLM for each state. The latent state evolves across trials according to a transition matrix that generally encourages state continuity across trials. **(d)** Normalized heldout log-likelihood (in units of bits per trial; see Methods) increased with the number of states; we selected the three-state model, where performance increases started to become minimal, to balance predictive performance with statistical power and interpretability of subsequent results. **(e)** State-specific GLM weights for the four model covariates: stimulus, previous choice (PC), win-stay/lose-switch (WSLS), and bias. The engaged state shows the strongest stimulus weight, whereas the biased states have strong opposing bias weights (right is a positive weight). **(f)** Empirical psychometric curves showing the probability of choosing right, as a function of the net amount of rightward sensory evidence. These were computed separately for trials in which the animal was in each inferred GLM-HMM state, illustrating differences in stimulus sensitivity and choice bias. **(g)** Percentage of trials assigned to each state. The engaged state accounts for approximately half of all trials, about twice the fraction assigned to either biased state. **(h)** The duration (mean ± s.e.m.) that animals remained within a given state prior to switching states, demonstrating persistent behavioral strategies.

### 2.1 A GLM-HMM identifies persistent behavioral strategies across datasets

To identify the behavioral strategies mice used during decision making, we modeled their choices with a GLM-HMM, which allows the relationship between task covariates and choice to vary across latent states (Fig. 1c). Here, as task covariates, we included the stimulus (the number of rightward minus leftward cues, z-scored), the previous trial’s choice (rightward or leftward), a win-stay/lose-switch term (the previous choice weighted by whether it was rewarded, so that the animal is predicted to repeat the previous choice after a reward and to switch after an error), and a bias (a constant offset reflecting a preference for rightward or leftward choices). We fit the GLM-HMM to choices on non-perturbation trials pooled across the three datasets comprising 54 mice, yielding a common set of states for examining how cortical inactivation affects behavior. The model allows the influence of task variables on choice to differ across discrete latent states while describing how those states persist or change across trials (Fig. 1b,c).

To select the number of latent states, we performed five-fold cross-validation and compared held-out log-likelihood (LL) across models with different numbers of states. Log-likelihood increased with model complexity, but the gains beyond three states were comparatively small (Fig. 1d), and we thus used a three-state model for subsequent analyses. We also note that the four-state model contained three almost-identical states to our selected three-state model (described below), with a rarely used (< 8% occupancy) fourth state. In our three-state model, in one state, choices depended strongly on the stimulus, consistent with an engaged strategy (Fig. 1e,f; red). The other two states (green and purple) showed opposing bias weights and weaker stimulus dependence, consistent with right- and left-biased strategies. The biased states nevertheless retained some sensitivity to the stimulus. In this framework, the engaged state accounted for approximately half of all trials, roughly twice the fraction assigned to either biased state (Fig. 1g). All states occurred within all datasets (Fig. S1c). Additionally, the mice persisted within inferred states for many trials—over 25 trials on average for biased states, and over 35 trials on average within the engaged state (Fig. 1h; Fig. S1a for example posteriors). Together, these results show that the relationship between different covariates and choice changes across persistent behavioral states. We next asked whether cortical inactivation affects decisions differently in each state.

### 2.2 Whole-trial cortical inactivation has state-dependent effects on decision making

We first examined the effect of whole-trial optogenetic inactivation on the behavior of 20 mice expressing the excitatory opsin Channelrhodopsin-2 in inhibitory interneurons (VGAT-ChR2-EYFP), whose activation with blue light led to reversible cortical silencing. On interspersed trials, one of 29 different bilateral cortical locations spanning the dorsal cortex was optogenetically inactivated, each with a probability of 0.01 (Fig. 2a). We focused on whether optogenetic effects depended on the animals’ underlying behavioral strategies. To assign states without using choices made under perturbation (‘opto’ trials), we fit the GLM-HMM as described above to control (‘non-opto’) trials and assigned each opto trial the state of its temporally nearest non-opto trial (Fig. 2b; Methods). Importantly, animals remained in the same behavioral state for many non-opto trials spanning many interspersed opto trials (Fig. 2c; arrows on top show interspersed opto trials). This means that the vast majority of opto trials occurred within the time period of an underlying broader behavioral state.

**Fig. 2:**
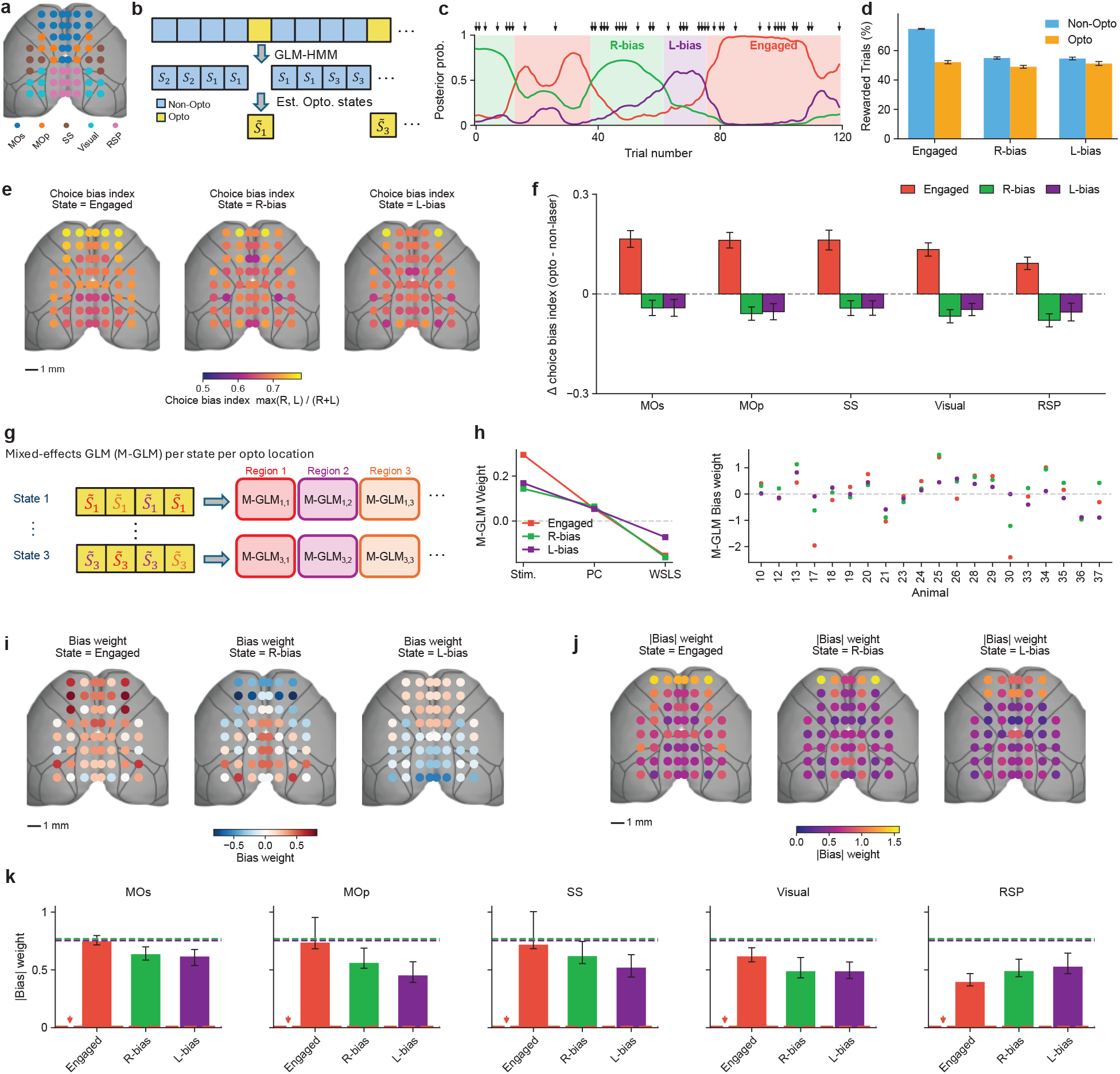
Whole-trial cortical inactivation alters choice bias in a state- and region-dependent manner. **(a)** 29 bilateral locations of whole-trial optogenetic inactivations. Locations are color-coded according to the grouping of anatomical regions that are used in subsequent panels for quantification. **(b)** Schematic of assigning opto trials to states. A GLM-HMM was fit to non-opto trials to infer behavioral states (*S*_*i*_ denotes state *i*). Opto trials were assigned the state of the nearest non-opto trial 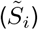. **(c)** Example posterior probabilities of the three behavioral states across trials, and the assigned trial per state. Arrows mark opto trials, demonstrating that these opto trials occur within persistent blocks of non-opto strategies. **(d)** Percentage of rewarded trials with and without inactivation, separated by behavioral state. Error bars are s.e.m. **(e)** Choice bias index across inactivation sites for each behavioral state, showing higher bias after frontal than posterior inactivation in the engaged state. **(f)** Change in choice bias index relative to non-opto trials across five cortical regions defined in panel a. Bias increases most strongly in the engaged state, particularly following frontal inactivation, whereas inactivation reduces bias in both biased states across all five groups of regions. **(g)** To quantify how inactivation affected choices within each state, a separate mixed-effects GLM model was fit to the corresponding opto trials in that state, separately for each inactivated region; colors denote different regions. **(h)** Example fixed-effect GLM weights for the three states (left) and animal-specific changes in the mixed-effect bias weight (right), allowing the model to capture variation in inactivation-induced bias across animals. **(i**,**j)** Cortical maps of signed bias weights and absolute bias weights, respectively, for each state. **(k)** Mean absolute bias weight across five cortical region groupings, separated by state. Dashed lines show the non-opto baseline. The resulting bias is greater in the engaged state after frontal inactivation, unlike in RSP.

Our previous analysis of this dataset, which had excluded low-performance periods, had revealed widespread effects of inactivation on behavior [18]. Extending these findings, here we observed that cortical inactivation impacted behavior across behavioral states. First, it significantly reduced the percentage of rewarded (successful) trials in all three behavioral states (Fig. 2d, p = 3.9 *×* 10^*−*10^, 2-way repeated-measures ANOVA). Reduced reward rates were observed across many cortical inactivation sites rather than at a small number of isolated locations, particularly in the engaged state (Fig. S2c). Moreover, investigating psychometric curves pooled across animals and for individual mice suggested that these decreases in reward rates relate to a reduced dependence on the stimulus across behavioral states, and may also relate to changes in levels of side bias, particularly in the engaged state (Fig. S2a,b).

Therefore, to explore the impacts of perturbation on the animals’ choice bias, for each inactivation site, we quantified its magnitude using the choice bias index, which ranges from 0.5 (across trials, an animal choosing half right and half left) to 1.0 (an animal chooses only right or only left). We did so separately for the engaged, right-biased, and left-biased states. By eye, it appeared that, in the engaged state, frontal inactivation led to the largest choice bias, relative to posterior inactivation (Fig. 2e, Fig. S2d). Additionally, the choice bias index appeared slightly larger following frontal inactivation in the engaged state, relative to the biased ones. To account for differences in baseline bias between states, we subtracted each state’s choice bias index computed on non-opto trials from its index on opto trials and summarized the effects across five cortical regional groupings (Fig. 2f). Inactivation increased bias in the engaged state, with an approximate gradient from weaker in posterior to stronger more frontally. Differences were significant between frontal regions and RSP (p = 0.017, paired t test with false discovery rate—FDR—correction, *n* = 20 mice). The direction of the effect reversed in the two biased states: inactivation reduced their pre-existing choice bias across all five regions, with a slightly larger (but not significant) observed reduction in the retrosplenial cortex (RSP). This suggests a striking state-dependent effect where inactivating a given location can increase or decrease behavioral bias depending on the animal’s decision-making strategy.

To more rigorously quantify the animals’ bias while accounting for correlations with other covariates and also bias variation across animals, we fit mixed-effects GLMs to the behavior on *inactivation* trials, separately for each behavioral state, and separately for each inactivation location (Fig. 2g; example fit for one location in Fig. 2h). In these models, stimulus, previous choice, and win-stay/lose-switch (WSLS) had weights shared across animals, whereas the bias term varied by animal, allowing the model to capture differences in the magnitude and direction of inactivation-induced bias between mice.

Investigating the learned model weights confirmed and extended the location- and state-dependent effects on bias seen with the choice bias index (Fig. 2h–k; also see Fig. S3 for the non-bias model covariates). In the engaged state, bias was indeed stronger in frontal regions. Interestingly, we also observed that frontal inactivation led to a reversal of side bias in biased states (Fig. 2i). That is, when animals were in a right-biased state, frontal inactivation commonly led to a leftward bias, and the reverse in the left-biased state. To further quantify the effect of silencing on bias, we grouped the absolute weight of the bias (to show strength of the bias across all animals, without concern for directionality, which could be inconsistent across animals) across the 5 larger anatomical groupings (Fig. 2k). Absolute bias became larger in engaged states versus biased states in frontal areas (p=0.02 in both frontal regions, comparing Engaged to average of R-bias and L-bias, FDR-corrected bootstrap test - see Methods). This corresponds to an interesting paradoxical effect where the bias increases in the engaged state relative to its non-opto baseline (red dashed line), while the bias actually decreases relative to its non-opto baseline in biased states (purple/green dashed lines). We again saw that the impact of optogenetic silencing on absolute bias in the engaged state was region-dependent, yielding much stronger bias in more frontal regions (MOs, MOp) than in posterior, in particular the RSP. Statistically, using a repeated measures ANOVA with factors behavioral state and region grouping, we saw a significant effect across the interaction of state and region (p = 0.008, *n* = 20). In total, these inactivation effects are suggestive of frontal areas being particularly involved in suppressing bias—as their inactivation leads to particularly strong bias.

### 2.3 The effect of inactivation depends on its timing within a trial

To provide greater understanding of the temporal evolution of the cortical contribution to behavioral strategies, we next analyzed sub-trial cortical inactivations. We focused on inactivations that occurred during the first and second halves of the cue region (0–100 and 100–200 cm; which we refer to as cue-Half1 and cueHalf2), and during the cue-free delay period preceding the turn (200–300 cm; Fig. 3a,b), enabling us to compare perturbations at different stages of the decision. Across the entire dataset, individual animals had different perturbation locations and epochs. We chose to focus on a subset of 15 mice (751 total sessions) that had bilateral perturbations in both anterior MOs (aMOs) and RSP, to enable a continued focus on the frontal versus posterior differences observed during whole-trial inactivations.

**Fig. 3:**
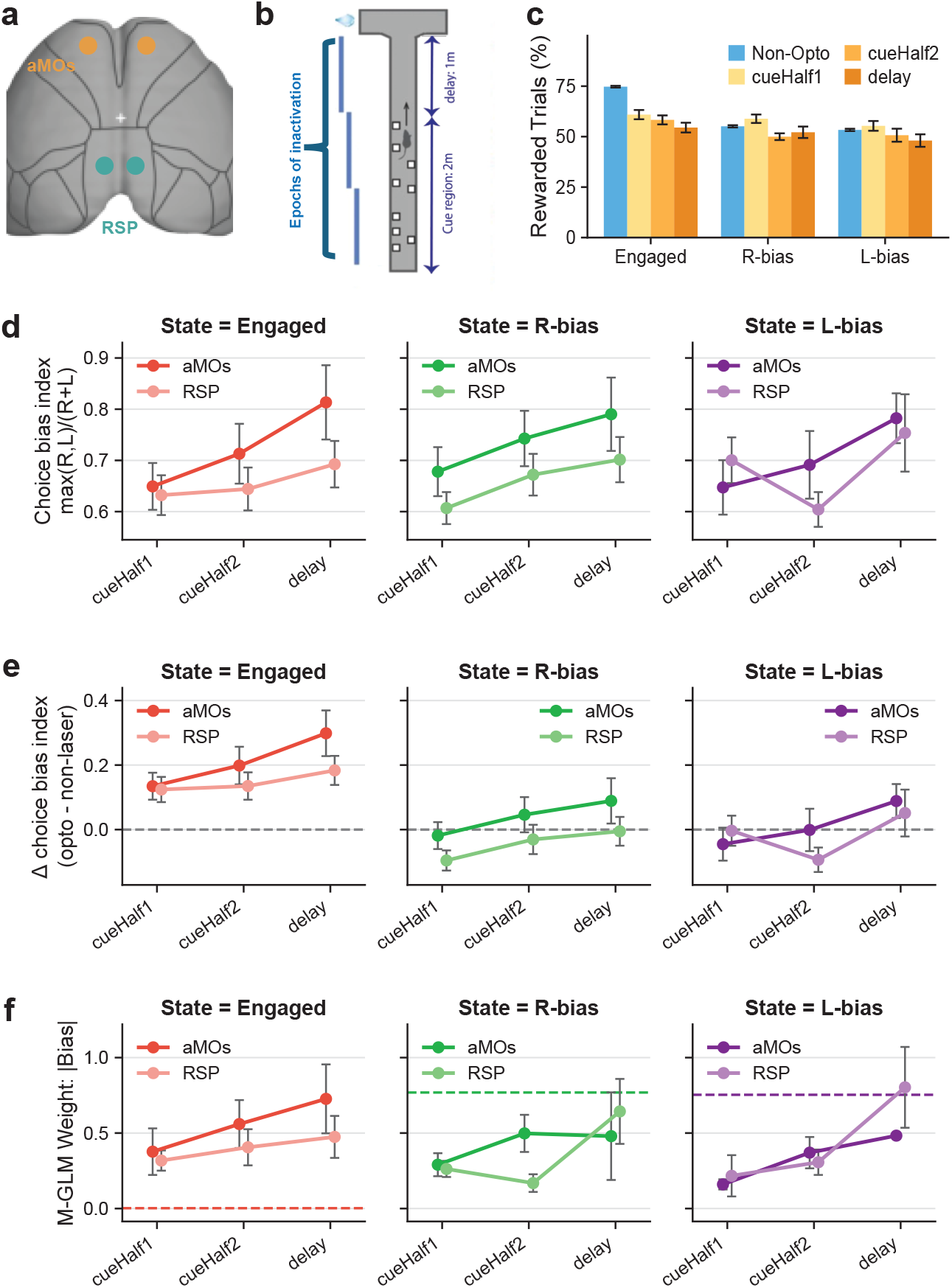
The effect of sub-trial cortical inactivation on choice bias depends on behavioral state, region, and task epoch. **(a)** Location of the two optogenetic locations focused on for sub-trial inactivation analyses. **(b)** Schematic of bilateral inactivation during selected portions of the cue region or the subsequent delay (memory) period. **(c)** Percentage of rewarded trials for non-opto trials and for inactivation during each of the three epochs. Inactivation has the largest effect on choice later in the trial, closer to the choice report. Error bars are s.e.m. **(d)** Choice bias index following aMOs or RSP inactivation during the three epochs, separated by behavioral state. Error bars are s.e.m. **(e)** Change in choice bias index relative to non-opto trials for the conditions in **(d). (f)** Absolute bias weights from state-specific mixed-effect GLMs for the same inactivation conditions; dashed lines indicate the baseline weights from non-opto trials.

As previously observed [19], inactivation reduced the percentage of rewarded trials more for inactivations occurring later in the trial (Fig. 3c, *p* = 2 *×* 10^*−*11^, *n* = 15, 2-way repeated-measures ANOVA with factors behavioral state and inactivation epoch). This was most noticeable in the engaged state, which had a higher non-opto reward rate (the biased states had just above 50% non-opto reward rates).

We next focused on the bias caused by inactivations, first by analyzing the animals’ choice bias index. As in the whole-trial inactivation, we observed that, during the engaged state, aMOs inactivation led to greater bias relative to RSP inactivation (Fig. 3d,e). Here, we observed that this effect was most prominent during the delay epoch. Relatedly, the effect of aMOs inactivation itself on bias increased later in the trial. Consistent with whole-trial inactivation results, the choice bias significantly increased relative to the non-opto baseline only within the engaged state (p < 0.001, two-sided one-sample t-test). This was true for both regions. It was also larger in aMOs, though the difference was not statistically significant.

To more rigorously isolate the impact of inactivations on choice bias, we fit mixed-effects GLMs (as in Fig. 2g) to inactivation trials, for each combination of region, state, and trial epoch (Fig. 3f; Fig. S4 for all parameter values). As with the choice bias index, during the engaged state, aMOs inactivation led to a greater but non-significant absolute bias weight relative to RSP inactivation (Fig. 3f). The absolute bias weight significantly increased relative to the non-opto baseline (dashed line) within the engaged state (p < 0.02, two-sided one-sample t-tests, for both regions), particularly during the delay, and decreased during the biased states—the same paradoxical effect we observed during whole-trial inactivations. Interestingly, here, the decreases in the biased states were more prominent during the cue-period inactivations. We also note that the flip in side-bias observed in the whole-trial inactivations was not observed in this dataset. Together, these results extend the whole-trial inactivation results—suggesting that frontal areas’ suppression of bias in engaged states is most important for behavior during the delay period, closest to decision report.

### 2.4 Mesoscale cortical activity depends on behavioral strategy

The state-dependent effects of cortical inactivation motivated us to investigate how cortical activity varied based on the animals’ underlying behavioral strategy. We thus analyzed widefield calcium imaging from 6 mice performing the task. We first investigated how the average activity within anatomically defined regions compared over the course of a trial, based on that trial’s underlying GLM-HMM behavioral state (Fig. 4a). In frontal areas like aMOs and medial MOs (mMOs), we observed that activity in the engaged state is slightly elevated relative to biased states throughout most of the trial, up until when the turn is being executed at the end of the delay region. This was generally not the case in more posterior regions like primary visual cortex (VISp) and RSP, whose average activity was not systematically different between engaged and biased states throughout the trial. The elevated activity in frontal areas during the engaged state may explain our previously observed larger optogenetic effects on frontal areas within the engaged state.

**Fig. 4:**
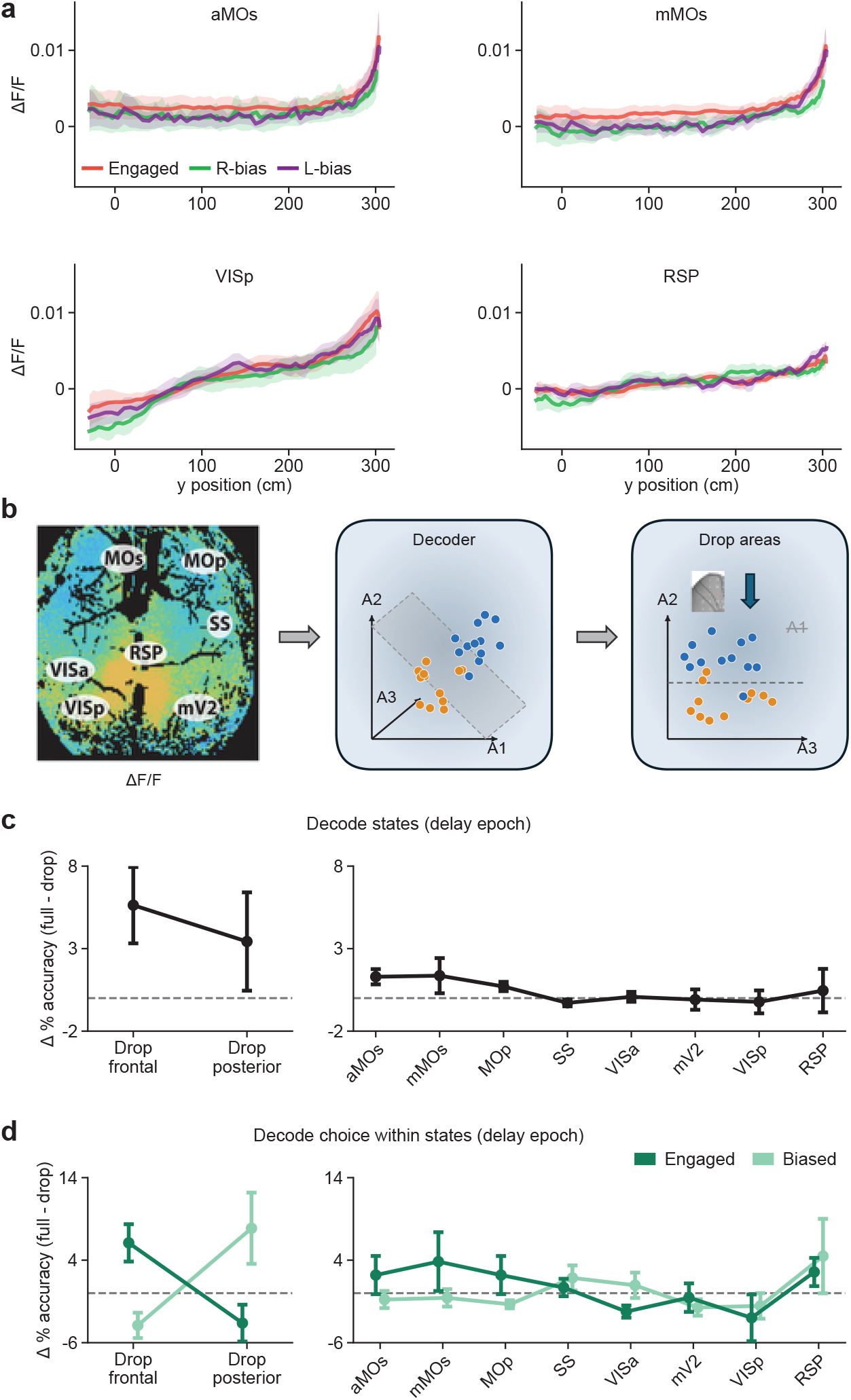
Frontal and posterior cortical activity make different contributions to decoding behavioral state and choice. **(a)** Mean neural activity (ΔF/F) along the maze in four cortical regions, separated by behavioral state. Frontal activity (aMOs, mMOs) is generally higher in the engaged than in the biased states over much of the maze. Shaded regions: s.e.m. across mice. **(b)** Analysis schematic. Decoders were fit to average widefield activity, with each region as a separate input, and one area or group of areas was excluded at a time to assess its contribution to decoding accuracy. **(c)** Decoding engaged versus biased states from delay-epoch activity (200– 300 cm). Loss of decoding accuracy (full minus reduced model, percentage points; mean ± s.e.m. across mice) after excluding the frontal (aMOs, mMOs, MOp) or posterior (VISa, RSP, mV2, VISp) regions together (left) or each region on its own (right). **(d)** Decoding choice within engaged and biased trials, using separate decoders and delay-epoch activity. Loss of choice-decoding accuracy after excluding frontal or posterior regions together (left) or each region on its own (right); mean ± s.e.m. across mice. Excluding frontal regions produced a positive loss in the engaged state, whereas in the biased state the largest losses came from posterior regions, in particular RSP, suggesting that the regional distribution of choice information depends on behavioral strategy.

To further quantify which cortical areas carried information about behavioral strategy, we assessed their contributions to behavioral-state decoding. We fit a logistic-regression classifier that predicted the trial’s behavioral state using the average neural activity across the trial’s delay epoch in each of 16 locations (two hemispheres for 8 regions). To determine the unique information about state in frontal areas (aMOs, mMOs, MOp), we removed these areas, refit the model, and calculated the decrease in accuracy using cross-validation (Fig. 4b). Indeed, frontal areas had significant unique information about the animals’ behavioral state (Fig. 4c; p = 0.030, *n* = 6, one-sided t test vs. zero). This effect was slightly greater, though not significantly so, relative to when posterior regions (VISa, mV2, VISp, RSP) were removed from the model. To more granularly view the particular regions that could have unique information about behavioral state, we took the same approach, removing one region (both hemispheres) at a time. In agreement with the more coarse analysis, it was primarily frontal regions that contributed positively towards behavioral state (Fig. 4c), consistent with the prominent state-dependent effects of frontal inactivation. Note, however, that the effect size for individual regions was substantially smaller (and not significant after multiple-comparisons correction), possibly related to information about behavioral strategy being somewhat redundant across frontal areas.

Next, given frontal regions’ most prominent optogenetic effects in the engaged state, we hypothesized that frontal areas may be particularly important for making decisions within that state. We thus tested the amount of information regions’ activity contained about choice within both the engaged and biased states. To do so, we took a comparable approach as above—we trained logistic-regression classifiers using all regions to predict the animal’s choice on each trial, and left regions out to gauge their unique information. Importantly, we trained separate classifier models for predicting choice within engaged and biased states (combining left-biased and right-biased states), enabling us to test whether regions were particularly important for encoding decisions within certain behavioral strategies. When first leaving out all frontal areas, we observed a substantial drop in choice decoding accuracy within the engaged state, but not the biased state (positive values in the engaged state; Fig. 4d, left). Strikingly, the effect reversed when dropping posterior areas from the model, in which choice decoding accuracy dropped within the biased state. This interaction between dropped area and behavioral state was statistically significant (p = 0.012, two-way repeated-measures ANOVA with factors of dropped area and state, *n* = 6). Moreover, these results were most prominent when using activity within the delay period (Fig. S5), compatible with the epoch-specific inactivations (Fig. 3). When holding out individual frontal regions from the model, we again saw confirmatory results (Fig. 4d, right, individual regions n.s.)—removing each of the three frontal areas from the model reduced performance (positive values in Fig. 4d) only within the engaged state and not the biased state. Results from removing individual posterior areas were not as clear, again potentially due to greater correlations between these areas yielding less region-unique information. Together, these results indeed suggest that frontal areas are particularly involved in making choices within the engaged state, while activity in more posterior areas is most related to decisions in biased strategies.

## 3 Discussion

We have applied GLM-HMMs to a large multi-modal dataset (Fig. 1) to discover that the causal relationship between activity and choice is reconfigured across the cortex depending on the behavioral strategy mice use to perform a decision-making task. Specifically, during optimal stimulus-driven (engaged) strategies, inactivation of distributed cortical regions decreases reliance on stimuli while increasing choice-side bias, with the largest effects from silencing frontal regions. Interestingly, the magnitude of behavioral effects from inactivations increases over the course of the trial for engaged strategies (Fig. 3). Finally, the results from the causal manipulations are supported by findings from mesoscale widefield calcium imaging data—i.e., that activity in frontal regions uniquely contribute to encoding choice during engaged, but not biased, strategies, with the opposite being true for posterior regions (Fig. 4). Our results extend recent findings from a few striatal and cortical regions [9, 10, 16] to show that strategy-dependent contributions to behavior are widespread.

While distributed, however, these causal contributions were not uniform across the cortex. Rather, frontal regions emerged as the most state dependent. Indeed, our findings from both imaging and inactivation are compatible with these regions being particularly important for generating sensory-evidence-based choices, and/or suppressing bias during sensory decisions. This is reminiscent of previous reports that prefrontal regions suppress impulsive actions during decision making [28, 29]. More generally, it is compatible with the fact that frontal regions, particularly MOs, are required for different perceptual decision-making tasks [2, 18, 19, 30–35] and encode decisions and multiple task strategies (e.g., [36–39]). Moreover, in a bandit task, MOs causally contributes to sub-optimal choice perseveration strategies, such that silencing it decreases side bias [40]. Our inactivation results from biased states are compatible with these recent findings, although, at least in the context of our task, MOs contributed most to engaged strategies.

Interestingly, posterior regions displayed the opposite pattern of strategy-dependent choice encoding, contributing most to this information in biased states, and with inactivation leading to the largest reduction of choice bias in those states in RSP. Although the latter trend was not statistically significant, these findings are compatible with recent reports implicating RSP and the posterior parietal cortex as integrators of trial history [5, 7, 41, 42]. Further, the dissociation in strategy-dependent activity patterns between posterior and frontal regions generally matches results from a recent report on internally generated motor strategies for solving a tactile decision-making task [17].

Our discoveries highlight the importance of using GLM-HMMs to infer latent behavioral states, given that animals (and mice in particular) use variable behavioral strategies to solve the same given task. Further, our combining multiple datasets under the same model gave us the statistical power and the ability to directly compare different evidence streams to understand the diverse state-dependent contributions different cortical regions have to decisions. While this approach assumed that comparable states happened across datasets, we did observe that the proportion of each state occurring in each dataset was fairly similar (Fig. S1c). One novel aspect of our approach, which enabled us to effectively integrate findings across opto and non-opto (observational) trials, was that we assumed that opto trials can be assigned to their nearest state defined from non-opto trials. While this assumption is based on the overall persistence of behavioral strategy across trials that we and others have observed, this of course, does not allow for the possibility that the inactivation itself induces a brief state switch. Still, the differences in the effects of optogenetic silencing based on the persistent non-opto states demonstrate that these more persistent (non-opto) strategies indeed impact the effects of opto.

An important remaining question is whether cortical activity sets the strategy and/or primarily reflects a strategy determined elsewhere. More generally, future studies should focus on the mechanisms that switch between and maintain different internally generated behavioral strategies. One possibility is that these mechanisms are shared with the ones implementing (externally imposed) task switches, in which frontal areas have been implicated [43–46]. Instead or additionally, neuromodulators such as acetyl-choline and norepinephrine have been proposed to facilitate cognitive flexibility [47, 48]. For instance, norepinephrine is important for switching in response to hidden changes in task contingencies [49–51] and cholinergic projections to the cortex have strategy-dependent dynamics during the accumulating-towers task [15].

Although many important questions remain, our study provides an important step in understanding context-dependent computation in the cortex, by revealing a striking reconfiguration of dynamics leading to choice by an internally generated behavioral strategy.

## 4 Acknowledgments

The datasets were collected in the laboratories of David W. Tank and Carlos D. Brody, funded by the NIH grant U19NS132720. Work was additionally funded by NIH grants K99/R00MH120047 (L.P.), R01MH138285 (L.P.), and R00 NS119787 (J.G.), and a McKnight Neuroscience Scholar Award (L.P.). Additionally, J.G. and L.P. acknowledge support from the National Institute for Theory and Mathematics in Biology through the National Science Foundation (DMS-2235451) and the Simons Foundation (MPTMPS-00005320). J.W.P was supported by grants from the Simons Collaboration on the Global Brain (SCGB AWD543027), the NIH BRAIN initiative (R01EB026946),and U19 NIH-NINDS BRAIN Initiative Awards (U19NS104648, U19NS123716).

## 5 Methods

### 5.1 Datasets and summary of experimental methods

This study combined three published experimental datasets as detailed below, with all procedures approved by the Institutional Animal Care and Use Committee at Princeton University and performed in accordance with the Guide for the Care and Use of Laboratory Animals. All mice were kept under a reversed 12/12 h light/dark cycle and underwent a single survival surgery procedure to render the skull over the dorsal cortex optically transparent and implant a titanium headplate [18]. After surgical recovery, the mice were water restricted and underwent behavioral training in the accumulating-towers task following a software-automated curriculum described elsewhere [6].

The task happened in a virtual-reality apparatus. Briefly, head-fixed mice ran on a Styrofoam® ball floating on compressed air. Ball displacements were detected with optical velocity sensors and yoked to movement through a 3-m virtual T-maze, projected onto a large toroidal screen. During the first 2 m, salient visual cues (towers) were displayed transiently along either maze wall—appearing when the mice were 10 cm away and disappearing 200 ms later. This was followed by a 1-m delay region with neutral wallpaper, after which the mice turned into one of the side arms, receiving a 10% sweet-condensed-milk reward if the chosen arm matched the side with highest tower count. The tower positions and counts were randomly drawn in each trial from spatial Poisson processes of different means on the rewarded and unrewarded sides (7.7 and 2.3/m, respectively). Further details on the apparatus and task, including the training curriculum, can be found elsewhere [6, 18, 19].

Below we provide a description of the datasets with their respective summarized methods (note that some mice contributed data to both whole-trial and sub-trial inactivation datasets, such that we analyzed data from a total of 54 mice). Refer to the original reports for further details on experimental and data-preprocessing methods [18, 19].

- Whole-trial inactivation data [18] (Figs. 1, 2, S1–S3). We used data from 20 VGAT-ChR2-EYFP mice of both sexes, aged 2–16 months (Jackson Laboratories stock no. 014548), from a total of 178,430 trials and 894 sessions. Briefly, for each mouse, we defined a grid of 29 bilateral locations based on stereotaxic coordinates from bregma (spanning from -3.50 to 3.50 mm AP, and 0.25 to 2.00 mm laterally). We ensured locations matched across sessions by using rigid image registration algorithms relative to a reference image taken on the first day of experiments. We used a custom-built laser-scanning system for inactivation. A blue (473-nm) laser was launched into a single-mode fiber optic, scanned with 2-D galvanometers and passed through an F-theta scan lens before reaching the cortex through the intact clear skull. We used a power of 6 mW and a square wave of 40 Hz (0.8 duty cycle), with a 100-ms ramp down to minimize activity rebound. Bilateral inactivation was achieved by switching the galvanometers between hemispheres at 200 Hz. We confirmed with electrophysiology that we silenced the cortical locations by exciting inhibitory interneurons expressing ChR2. Inactivation onset and offset were yoked to the animals’ position in the maze, happening between the start of the cue region and the end of the delay region. Each bilateral location was inactivated with a probability of 0.01 (overall inactivation probability of 0.29). Grid locations were assigned to an anatomically defined region from the Allen atlas (common coordinate framework 3.0, CCF3.0) via a custom registration algorithm. Areas were then grouped into primary and secondary motor cortices (MOp and MOs), all subdivisions of the somatosensory cortex (SS), all subdivisions of the visual cortex within the inactivation grid (VISp, VISam, VISpm, VISa; collectively referred to as Visual), and granular and agranular retrosplenial cortex (collectively referred to as RSP).
- Sub-trial inactivation data [19] (Figs. 3, S1, S4). We used data from 28 VGAT-ChR2-EYFP mice of both sexes, aged 2–16 months, from a total of 239,351 trials and 1,194 sessions. We used the same methods as above for inactivation. The experiments targeted nine sets of bilateral cortical areas during six behavioral epochs defined by maze position: 0–50 cm (‘CueQuart1’), 50–100 cm (‘CueQuart2’), 100–150 cm (‘CueQuart3’), 0–100 cm (‘cueHalf1’), 100–200 cm (‘cueHalf2’), 200–300 cm (‘delay’). Inactivation of each area set happened with a probability of 0.05–0.15. Unlike the whole-trial dataset, different mice in this one experienced different subsets of the 54 experimental conditions. Thus, although all the control trials from this dataset were used to train the combined GLM-HMM, we focused the inactivation analysis on two regions for which we had enough mice who experienced both, specifically in the ‘cueHalf1’, ‘cueHalf2’, and ‘delay’ epochs (n = 15): the anterior part of MOs (aMOs) and the retrosplenial cortex (RSP). The stereotaxic coordinates for these two regions were, respectively: +3.0 mm AP, 1.0 mm ML and –2.5 mm AP, 0.5 mm ML.
- Widefield calcium imaging data [18] (Figs. 4, S1, S5). We used data from 6 mice from a triple transgenic cross expressing the calcium indicator GCaMP6f in cortical excitatory neurons, of both sexes, aged 2-8 months (Ai93-D;CaMKIIa-tTA, Jackson Laboratories stock no. 024108 Emx1-IRES-Cre stock no. 005628). We had a total of 5,321 trials and 25 sessions. We used a custom-built widefield macroscope with tandem 1 *×* and 0.63 *×* lenses and alternating blue (470 nm) and violet (405 nm) excitation for isosbestic correction of hemodynamic contamination, with an effective frame rate of 10 Hz and field of view of approximately 10 *×* 10 mm. Per-area ΔF/F was computed by averaging raw fluorescence from each excitation channel across pixels within an area and applying a regression-based hemodynamic correction algorithm. Cortical area assignments are based on registering the widefield images to the Allen CCF3.0 using fiducial markers from both visual-field maps and anatomical features. We grouped subdivisions into the following eight areas, separately for each hemisphere: anterior and medial MOs (aMOs, mMOs), MOp, RSP, SS, VISa (or posterior parietal cortex, PPC), mV2 (VISam and VISpm combined), VISp. MOs was divided into the subdivisions by a line running anteroposteriorly at 1.25 mm ML. Activity was aligned to behavior and downsampled to 10-cm position bins by selecting the ΔF/F value from only the first frame after the animal crossed a certain y position in the maze.

### 5.2 GLM-HMM Overview

A Hidden Markov Model (HMM) describes sequential data as generated by a system evolving through unobserved, discrete latent states 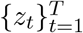, and has been widely used in neuroscience, computational biology, and behavioral analysis [52–55]. In an HMM, state transitions follow a discrete-time Markov process governed by a transition matrix *A* ∈ ℝ^*K×K*^ , with *P* (*z*_*t*+1_ = *k* | *z*_*t*_ = *j*) = *A*_*jk*_. Within a GLM-HMM, each discrete state has a state-specific GLM observation model (described below). This GLM-HMM architecture lets the influence of task covariates on choice vary flexibly with latent behavioral strategy [9, 12, 22–25, 56, 57].

#### Observation model

Conditioned on latent state *z*_*t*_ = *k*, we modeled choice with a Bernoulli GLM: a response *y*_*t*_ = 1 denotes a rightward choice, and the probability of a rightward choice follows a logistic (logit-link) function of the trial covariates **x**_*t*_ (described below) and a state-specific weight vector **w**_*k*_:

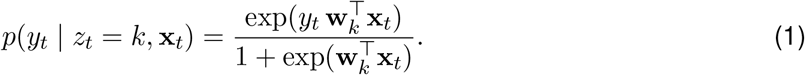

The weights **w**_*k*_ thus capture each state’s sensitivity to each covariate, letting us read off behavioral strategy directly from the fitted weights. We estimated **w**_*k*_ by maximum a posteriori (MAP) estimation under a zero-mean Gaussian prior, which penalizes large weights and improves generalization [58]. A detailed mathematical formulation of the GLM-HMM is found within Section 5.7, including full specifications of the prior. We assessed generalization with cross-validation on held-out sessions (Section 5.8).

#### Covariates of the model

Within this framework, we incorporated several task-relevant covariates into the GLM component of the model. These included: stimulus, previous choice (PC), win-stay/lose-switch (WSLS), and a bias term. Each of these covariates played a specific role in capturing different aspects of the animal’s behavior.

The “stimulus” was defined as the difference between the number of towers on the right and left sides of the maze on each trial. To ensure consistency across sessions, stimulus values were normalized by dividing by the across-session standard deviation of trial contrasts. “Previous choice” was encoded as a binary variable taking values of *−*1 or +1, indicating whether the mouse’s choice on the preceding trial was to the left or right, respectively. The “win-stay/lose-switch” (WSLS) covariate reflected behavioral adaptation to the previous trial’s outcome, computed as the previous choice (*−*1 left, +1 right) multiplied by the previous trial’s reward outcome (+1 rewarded, *−* 1 unrewarded). It therefore took the value +1 following either a rewarded rightward choice or an unrewarded leftward choice (both predicting a rightward choice on the current trial), and *−*1 following either a rewarded leftward choice or an unrewarded rightward choice (both predicting a leftward choice). A constant “bias” term was also included to capture any inherent side preferences independent of the other covariates.

#### Model Fitting

To recover latent behavioral states, we applied the GLM-HMM to all three datasets by combining data across all animals into a single pooled dataset. This approach enabled us to obtain a global fit of the model, ensuring that the inferred states are consistent across datasets, animals, and trials, and that the resulting model captures the shared underlying structure of the behavior. Details about the model fitting procedure can be found in Section 5.7.

### 5.3 Quantifying psychometric curves

To quantify the animal’s stimulus-driven behavior, we computed psychometric curves by plotting the probability of a rightward choice as a function of net sensory evidence, defined as the number of right minus left cues. Net evidence (range *−*15 to +15 cues) was divided into six equal-width bins. For each bin we computed the fraction of trials with a rightward choice, 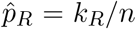, where *k*_*R*_ is the number of rightward choices and *n* the number of trials in the bin. In Fig. 1f, curves were computed on non-opto trials, pooled across animals and separated by the GLM-HMM state assigned to each trial; Fig. S2a,b shows the same analysis for non-opto and opto trials, pooled across animals (a) and for an example mouse (b). These curves are empirical; no parametric function was fit. Their slope reflects stimulus sensitivity and their horizontal offset reflects side bias.

### 5.4 Determining optogenetic effects on behavior based on underlying states

To identify behavioral states in the optogenetic (opto) trials, we first fit a GLM-HMM to the non-optogenetic (non-opto) data in order to estimate the animal’s latent state on each trial, as described in previous sections. These state assignments served as a reference for the opto trials. For each opto trial, we assigned a state by selecting the temporally nearest non-opto trial and using its inferred state label. This allowed us to align each opto trial with an estimated behavioral state derived from the animal’s typical (non-opto) behavior.

For each inactivation site and behavioral state, we additionally computed a choice bias index, defined as CBI = max(*p*_*R*_, *p*_*L*_)*/*(*p*_*R*_ +*p*_*L*_), where *p*_*R*_ and *p*_*L*_ denote the fractions of rightward and leftward choices, respectively; this index ranges from 0.5 (equal numbers of left and right choices) to 1.0 (choices entirely to one side).

We then examined how optogenetic inactivation impacted behavior within each of these inferred states by analyzing reward rates and state-specific choice biases. To formally quantify the impact of inactivation on behavioral strategy, we fit separate mixed-effect GLMs to the opto data for each inferred state, as described in the next section. These per-state models were applied either across brain regions or— when relevant—across region-by-epoch combinations, enabling us to characterize how task covariates influenced decisions under inactivation, conditioned on the animal’s behavioral state.

### 5.5 Mixed-effects GLM

To quantify how cortical inactivation affected choice within each behavioral state while accounting for differences in side preference across animals, we used a mixed-effects generalized linear model (M-GLM). This model extends the Bernoulli GLM observation model described above by allowing each animal to have its own intercept (bias) term, while sharing the stimulus, previous-choice, and win-stay/lose-switch (WSLS) coefficients across animals. Separate M-GLMs were fit to optogenetic trials within each inferred behavioral state and inactivation location, or within each state and region–epoch combination for the subtrial inactivation dataset.

For a binary choice *y*_*t*_ *∈ {*0, 1*}*, with 1 denoting a rightward choice, the Bernoulli GLM relates the probability *p*_*t*_ of a rightward choice to a linear predictor *η*_*t*_ through the logit link, 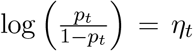 ,or equivalently 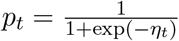. The probability of the observed choice is therefore

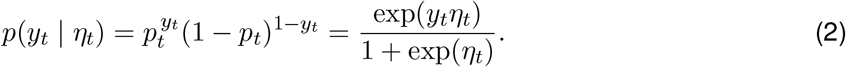

In the M-GLM, the linear predictor for a trial contributed by animal *a*(*t*) was

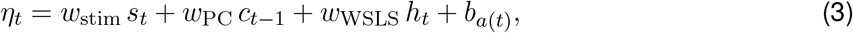

where *s*_*t*_, *c*_*t−*1_, and *h*_*t*_ denote the stimulus, previous-choice, and WSLS covariates, respectively, as defined in Section 5.2. The weights *w*_stim_, *w*_PC_, and *w*_WSLS_ were shared across animals within each fitted model, whereas *b*_*a*(*t*)_ captured the animal-specific side preference within the given behavioral state and inactivation condition. Sharing the stimulus and history coefficients pools information across animals, while animal-specific intercepts accommodate differences in side preference.

Equivalently, the per-animal biases can be expressed as coefficients on animal-indicator covariates. For the *M* animals contributing to a given fit, we defined

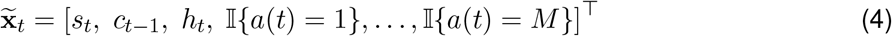

and

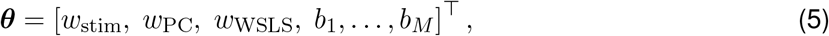

so that 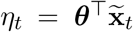. The *M* animal indicators replace a single shared constant bias term, and no additional common intercept was included.

Parameters were estimated by maximum likelihood. For the set of trials *T*_*m*_ contributed by animal *m* within the given state and inactivation condition, the log-likelihood was

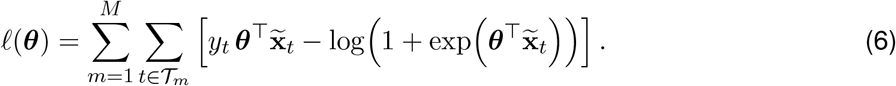

The fitted *b*_*m*_ quantify signed side preference after accounting for stimulus- and history-dependent contributions to choice: positive values favor rightward choices, whereas negative values favor leftward choices. Their absolute values were used to compare bias magnitude across behavioral states, cortical locations, and inactivation epochs (Figs. 2 and 3).

To determine statistical significance of differences in the bias weights between engaged and biased states (Fig. 2k), we resampled the inactivation trials with replacement (i.e., bootstrapped), separately for each state and region, and refit the mixed-effects GLM on each resample, 100 times. In each resample we computed the average absolute bias weight across animals. We compared the engaged state with the average of the right- and left-biased states in each region. The p-values come from the empirical distribution of that difference (two-sided), FDR-corrected across the five regions (Benjamini–Hochberg).

### 5.6 Decoding from neural activity

To test how choice- and strategy-related information is distributed across cortex, we trained two families of logistic-regression decoders on widefield activity from eight cortical areas (aMOs, mMOs, MOp, SS, VISa, mV2, VISp, RSP; 16 hemispheric ROIs), using cross-validation. We used L2-regularized logistic regression with standardized predictors and balanced class weights; the regularization strength, *C*, was selected using cross-validation. Imaging frames were assigned to maze position and activity was averaged within three position epochs (0–100, 100–200, 200–300 cm) or over all epochs. Session-level z-scoring was applied separately to neural activity before decoder fitting.

The first decoder predicted the trial’s behavioral state (Engaged vs. Biased, with R-bias and L-bias pooled) from delay-epoch (200–300 cm) activity across all 16 ROIs. To quantify each area’s contribution, we refit this decoder after excluding both hemispheres of the frontal group (aMOs, mMOs, MOp) or the posterior group (VISa, RSP, mV2, VISp) together, and separately after excluding one region (both hemispheres) at a time, defining the accuracy lost as the accuracy of the full decoder minus that of the reduced decoder (Fig. 4c).

The second decoder predicted the animal’s binary choice (left vs. right) from the same regional activity, fit separately within Engaged and within Biased trials (R-bias and L-bias pooled) to test whether regions carried choice information differentially by strategy. As with the state decoder, we quantified each region’s unique contribution using delay-epoch activity by excluding the frontal or posterior region groups together, and by excluding individual regions, and defined the accuracy lost analogously (Fig. 4d). The corresponding analysis using the first (0–100 cm) and second (100–200 cm) cue-region epochs, along with using average activity over all epochs, is reported in Fig. S5.

### 5.7 GLM-HMM Inference

#### 5.7.1 Overview and Priors

We estimate the parameters of the GLM-HMM using Maximum A Posteriori (MAP) inference. The set of model parameters is denoted by Θ = {*A*, **w, *π***} , where *A* is the state transition matrix, **w** contains the observation weights across all states, and ***π*** is the initial state distribution. These components will be described in further detail throughout this section.

MAP estimation provides a principled approach to parameter inference by combining observed data with prior knowledge. Rather than identifying the parameter set that maximizes the likelihood alone, MAP seeks the mode of the posterior distribution over parameters. This approach is particularly useful when working with limited or noisy data, or when incorporating structured prior assumptions into the model [58].

To estimate Θ, we adopt the Expectation-Maximization (EM) algorithm, originally introduced by Dempster et al. [59]. EM is a widely used iterative procedure for fitting models with latent variables, and it has been successfully applied in related settings, including HMMs with external regressors [20, 22, 60]. We describe the algorithm in detail in the subsequent section.

Each iteration of EM involves two steps: the E-step computes expectations over the latent variables given current parameter estimates, and the M-step updates parameters to maximize the expected complete-data log-likelihood. These steps repeat until convergence to locally optimal parameters. The initial state distribution ***π*** ∈ ℝ^*K*^ is drawn from a Dirichlet prior with parameter *γ*_*π*_ = 1. For the state-dependent observation weights **w**_*k*_, we place independent Gaussian priors with zero mean and variance *σ*^2^, applied separately to each state *k*. This prior structure follows the implementation in the ssm package [61], which we used to fit the GLM-HMM, and matches the approach used in prior GLM-HMM analyses of mouse decision-making [12, 13]. The overall prior over model parameters is *p*(Θ) *≡ p*(*A*) *p*(***π***) *p*(**w**), where the prior over weights and initial condition is defined as:

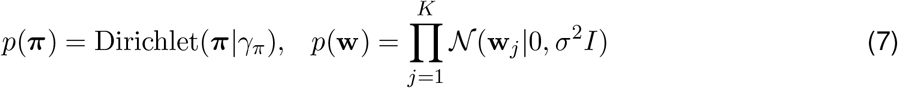

For the transition matrix *A*, we used a stationary transition model with a symmetric row-wise Dirichlet prior. Each row *A*_*j*_ defines a probability distribution over the next state. In the ssm parameterization, *α* controls the symmetric Dirichlet concentration, whereas *κ* adds prior mass specifically to self-transitions. We used *α* = 1 and *κ* = 0, corresponding to a uniform prior over each transition row with no additional prior preference for self-transitions:

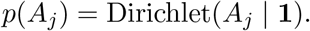

The transition matrix was initialized with high diagonal probabilities, but persistence in the fitted transition matrix was determined by the data.

The full set of observed inputs and outputs across trials is denoted by *F ≡ {***Y, X***}*, where **X** = *{x*_1_, … , *x*_*T*_ *}* are the trial covariates and **Y** = *{y*_1_, … , *y*_*T*_ *}* are the binary choices made by the mouse. In this setup, the Bernoulli GLM plays a critical role by mapping the input covariates to choice probabilities through the observation weights **w**_*k*_, allowing the model to capture the decision strategy within each latent state. The strength of the weight prior is tuned via grid search over different values of *σ*^2^.

Ultimately, the EM algorithm maximizes the log-posterior over model parameters, combining the data likelihood and the parameter priors. The resulting optimization objective is:

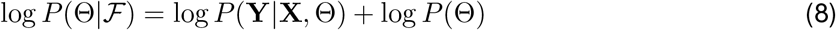

#### 5.7.2 Model initialization and fitting strategy

Because the expectation–maximization (EM) algorithm guarantees convergence only to a local optimum, the initialization of the GLM–HMM weights and transition matrix can substantially affect the fitted model. We therefore used a structured, multi-start initialization procedure.

First, we fit a one-state Bernoulli GLM to the pooled dataset using maximum-likelihood estimation, ignoring latent-state structure. This model provided an overall, state-averaged estimate of the relationship between the task covariates and choice. To initialize a *K*-state GLM–HMM, we duplicated this weight vector across all *K* latent states and added small, independent Gaussian perturbations to each copy. Thus, the state-specific GLMs were initialized near the one-state solution but were not identical, breaking symmetry and allowing the states to differentiate during fitting. The transition matrix was initialized to favor state persistence, with high probabilities on the diagonal and the remaining probability distributed uniformly across off-diagonal transitions, consistent with behavioral strategies persisting across consecutive trials.

Using this warm-start procedure, we fit the *K*-state GLM–HMM to the pooled dataset as described in Section 5.7.3, allowing each latent state to have its own GLM weights and transition probabilities. For each value of *K* and each cross-validation training fold, we performed many random initializations, with independent Gaussian perturbations applied to the initial GLM weights. Each model was fit using EM until convergence, and the initialization with the highest training log-likelihood was retained. This multistart procedure reduced sensitivity to poor local optima, a known challenge in EM-based optimization [62].

#### 5.7.3 Parameter estimation through the EM algorithm

We fit the GLM-HMM using the Expectation-Maximization (EM) algorithm [59, 63], a standard iterative procedure for maximum-likelihood or maximum-a-posteriori estimation in latent-variable probabilistic models such as HMMs [20]. The observed-data log-likelihood requires marginalizing over all latentstate sequences **Z**:

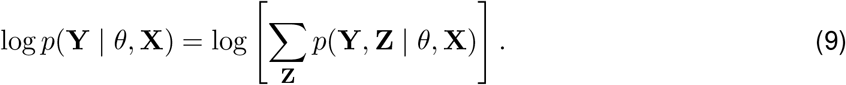

This sum contains *K*^*T*^ terms for a session of *T* trials, making explicit enumeration computationally prohibitive for long sessions. The forward algorithm computes the likelihood exactly, while forward– backward recursions compute the posterior state probabilities and expected transition counts needed for EM, with computational complexity *O*(*TK*^2^). EM alternates between two steps. In the E-step, we compute the posterior distribution over latent states given the current parameter estimates *θ*^(*i*)^, using the standard forward-backward algorithm for HMMs. The forward variable

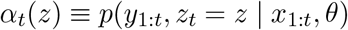

is the unnormalized joint probability of the observations up to trial *t* and the state at trial *t*. It was initialized as

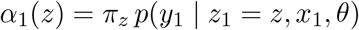

and updated recursively according to

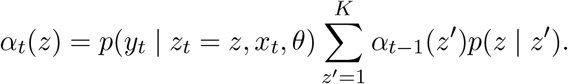

The backward variable *β*_*t*_(*z*) gives the probability of all future observations given the state at trial *t* and was computed using the corresponding backward recursion. Combining the two yields the singlestate posterior 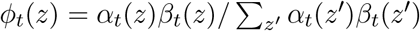 and the pairwise posterior over consecutive states *µ*_*t*_(*z, z*^*′*^), which together give the expected complete-data log-likelihood,

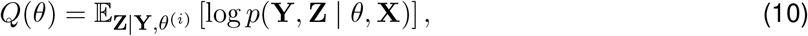

which decomposes into terms involving the GLM observation weights, the transition matrix, and the initial-state distribution. Because we performed MAP rather than maximum-likelihood estimation (Section 5.7.1), the M-step maximized *Q*(*θ*) + log *p*(*θ*) rather than *Q*(*θ*) alone, incorporating the Gaussian prior on the GLM weights and the Dirichlet priors on the transition matrix and initial-state distribution. Because of our uniform prior on the transition matrix, the transition-matrix MAP update reduces to row-normalizing the expected transition counts obtained from *µ*_*t*_(*z, z*^*′*^). The initial-state distribution *π* was updated analogously, as the (Dirichlet-regularized) average of the single-state posterior *ϕ*_1_(*z*) at the first trial of each session across all sessions. In the M-step, the GLM observation weights have no closed-form update because of the Bernoulli nonlinearity. The GLM–HMM was implemented using the ssm package. We maximized the relevant component of the expected complete-data log-posterior separately for each state using SciPy’s trust-region Newton conjugate-gradient algorithm (trust-ncg), with analytically derived gradients and Hessians supplied by the ssm observation model [64]. We alternated E- and M-steps until the change in the training objective between successive iterations fell below a fixed tolerance or the maximum number of iterations was reached. As described in Section 5.7.2, we fit each model from multiple initializations and retained the solution with the highest training objective.

### 5.8 GLM-HMM selection and predictive performance assessment

To assess the predictive accuracy and generalizability of the GLM-HMM, we employed a 5-fold crossvalidation procedure. The dataset was randomly partitioned such that approximately 80% of the sessions were allocated for training, while the remaining 20% were held out for testing. Importantly, sessions were sampled evenly across animals to ensure that inter-animal variability did not bias the evaluation. Model parameters were estimated exclusively using the training data via the Expectation-Maximization (EM) algorithm, while the held-out sessions were used solely for evaluating model performance.

To quantify model fit, we computed the log-likelihood of the test data for models with varying numbers of latent states (*K* = 1 to 5). For each test fold, the likelihood of the observed choices under the trained model was calculated using a single forward pass, without re-estimating parameters. For each held-out session *s ∈ S*_test_, let *T*_*s*_ be the number of trials in the session and *α*_*s,t*_(*z*) = *p*(*y*_*s*,1:*t*_, *z*_*s,t*_ = *z* | *x*_*s*,1:*t*_, *θ*) the unnormalized forward message at trial *t*, i.e. the joint probability of the choices up to trial *t* and of state *z* at trial *t*. The log-likelihood of session *s* was computed from the forward messages at the final trial of that session, 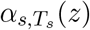:

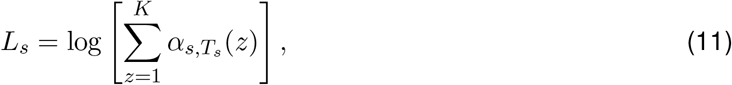

and the total held-out log-likelihood was

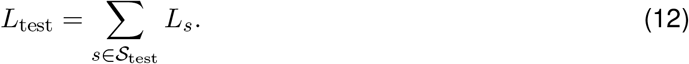

The recursion was evaluated in log space using the log-sum-exp operation for numerical stability. To facilitate comparison across models and datasets, we expressed the log-likelihood in units of bits per trial (bpt). This normalized measure captures how much better the GLM-HMM predicts test data compared to a baseline model that assumes independent Bernoulli trials. For each cross-validation fold, the baseline log-likelihood, *L*_0_, was computed for the held-out test data under a Bernoulli model whose rightward-choice probability was set to the empirical fraction of rightward choices trials in the corresponding training fold. The improvement in predictive performance was then computed as:

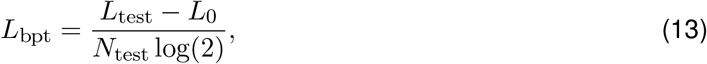

where *N*_test_ is the total number of included trials in the held-out test set. This formulation yields the normalized log-likelihood in bits per trial, which enables a clearer interpretation of model performance. A value of *L*_bpt_ *>* 0 indicates that the GLM-HMM outperforms the baseline model.

## 6 Data and code availability

Whole-trial optogenetic inactivation and widefield calcium imaging data analyzed in this study are partially available at https://data.mendeley.com/datasets/d2dkk9647b/1, and the dataset from the subtrial optogenetic inactivation experiments is available at https://doi.org/10.6084/m9.figshare. 19543948. The remainder of the raw data will be made available upon publication of this work. All code will be made available on Github upon publication of this work.

## 7 Author Contributions

ZM developed the model and performed the data analyses under the supervision of LP and JG. ZM, LP, and JG wrote the manuscript. All authors conceived the project, and revised and edited the manuscript.

## 9 Supplementary

**Fig. S1:**
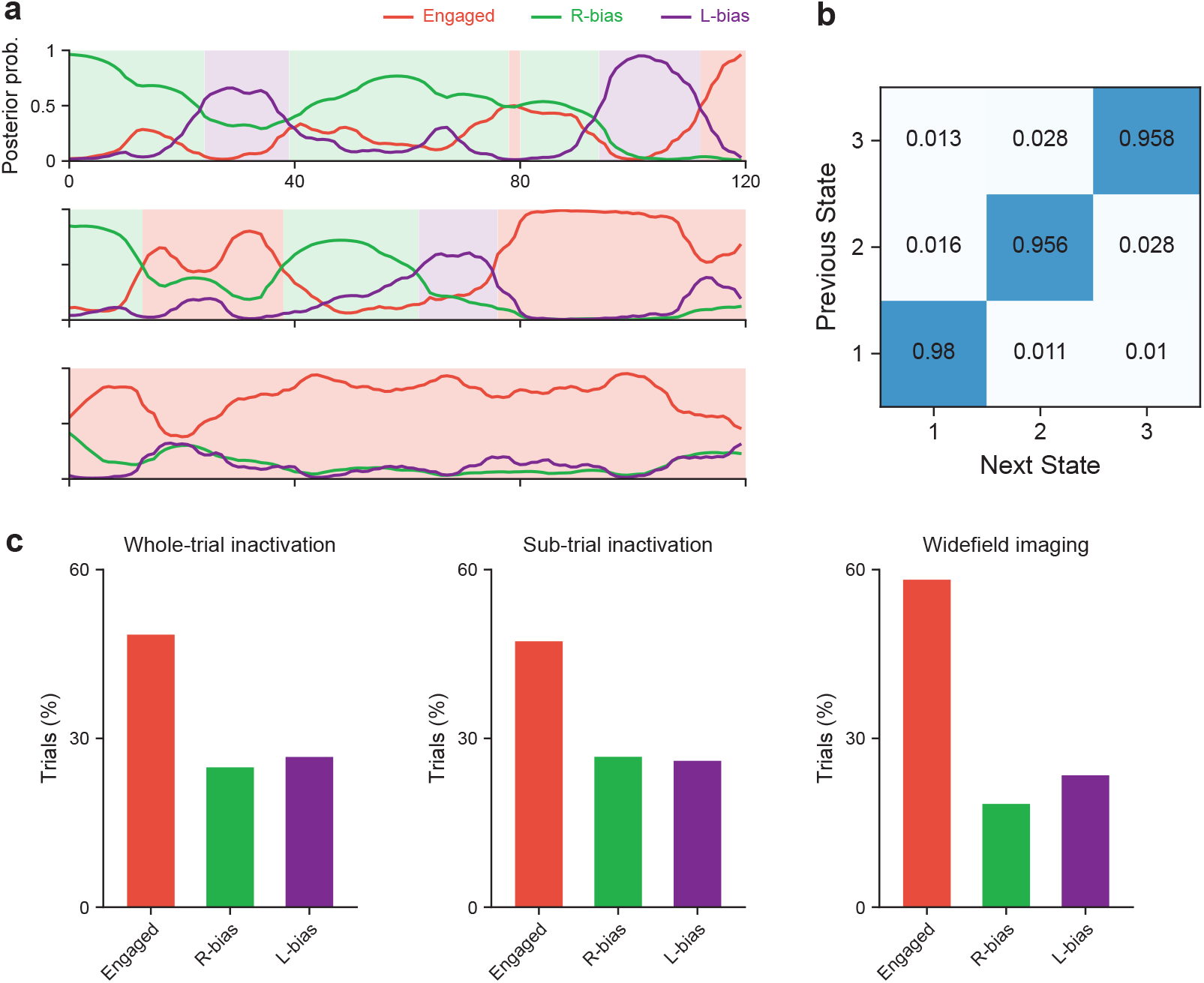
Example state posteriors, transition probabilities, and state occupancy of the GLM-HMM. **(a)** Posterior probability of each latent state over three example sessions from one mouse. Background shading marks the most probable state on each trial. **(b)** Transition probabilities between states (rows, previous state; columns, next state). High diagonal values show that states are persistent across trials. **(c)** Percentage of trials assigned to each state, shown separately for each of the three datasets. All three states occur in every dataset, and the Engaged state is the most common state in each.

**Fig. S2:**
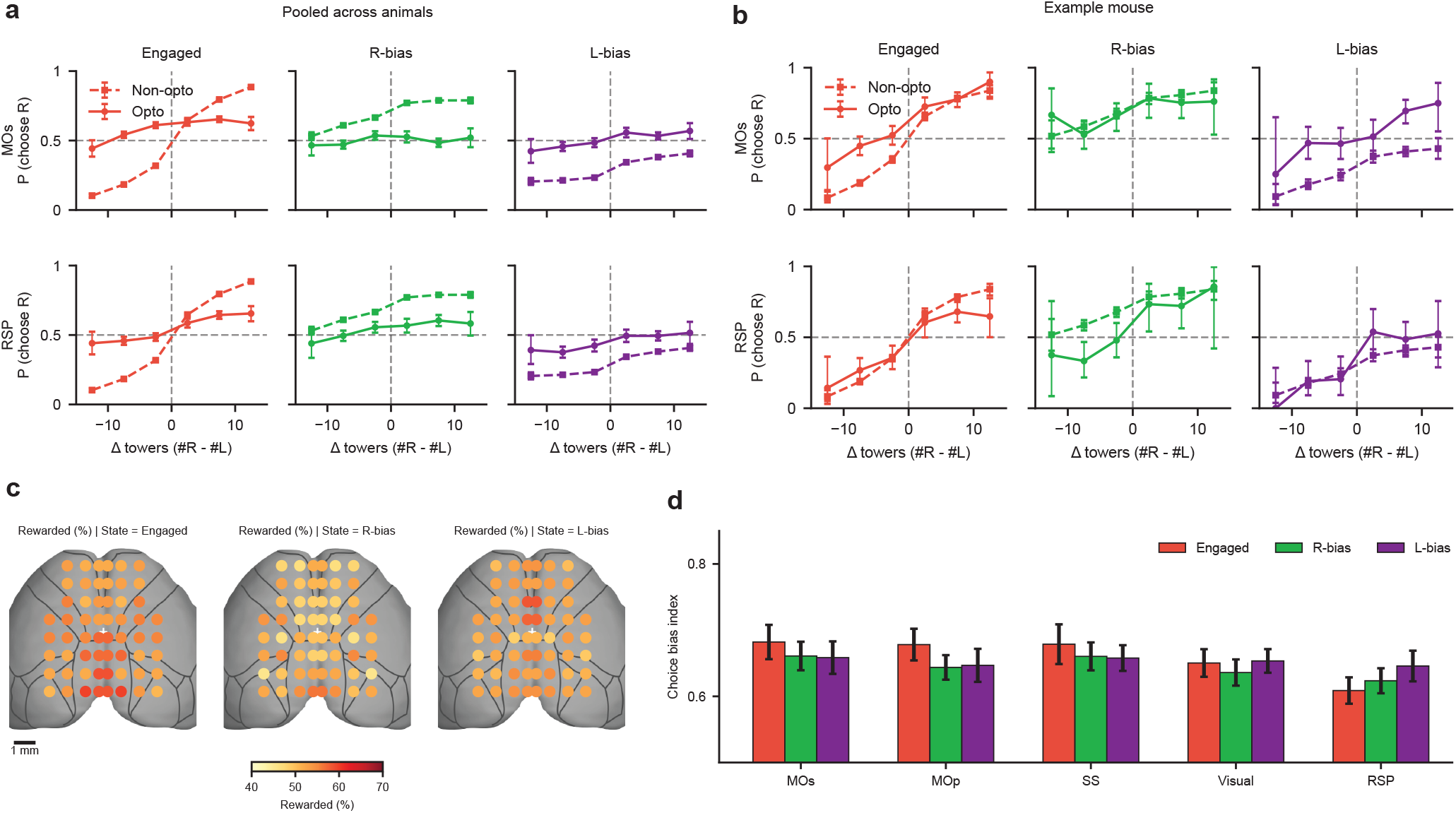
Psychometric performance and regional bias values following whole-trial cortical inactivation. **(a)**Psychometric curves, as in Fig. 1f, pooled across animals for non-opto and opto trials following MOs or RSP inactivation, separated by behavioral state. **(b)** Corresponding curves for an example mouse. **(c)** Percentage of rewarded trials across the 29 cortical inactivation sites, separately for each behavioral state. **(d)** Choice bias index across five anatomical groupings and the three behavioral states, showing slightly larger choice bias in engaged states relative to biased states after inactivation of frontal areas, but not RSP.

**Fig. S3:**
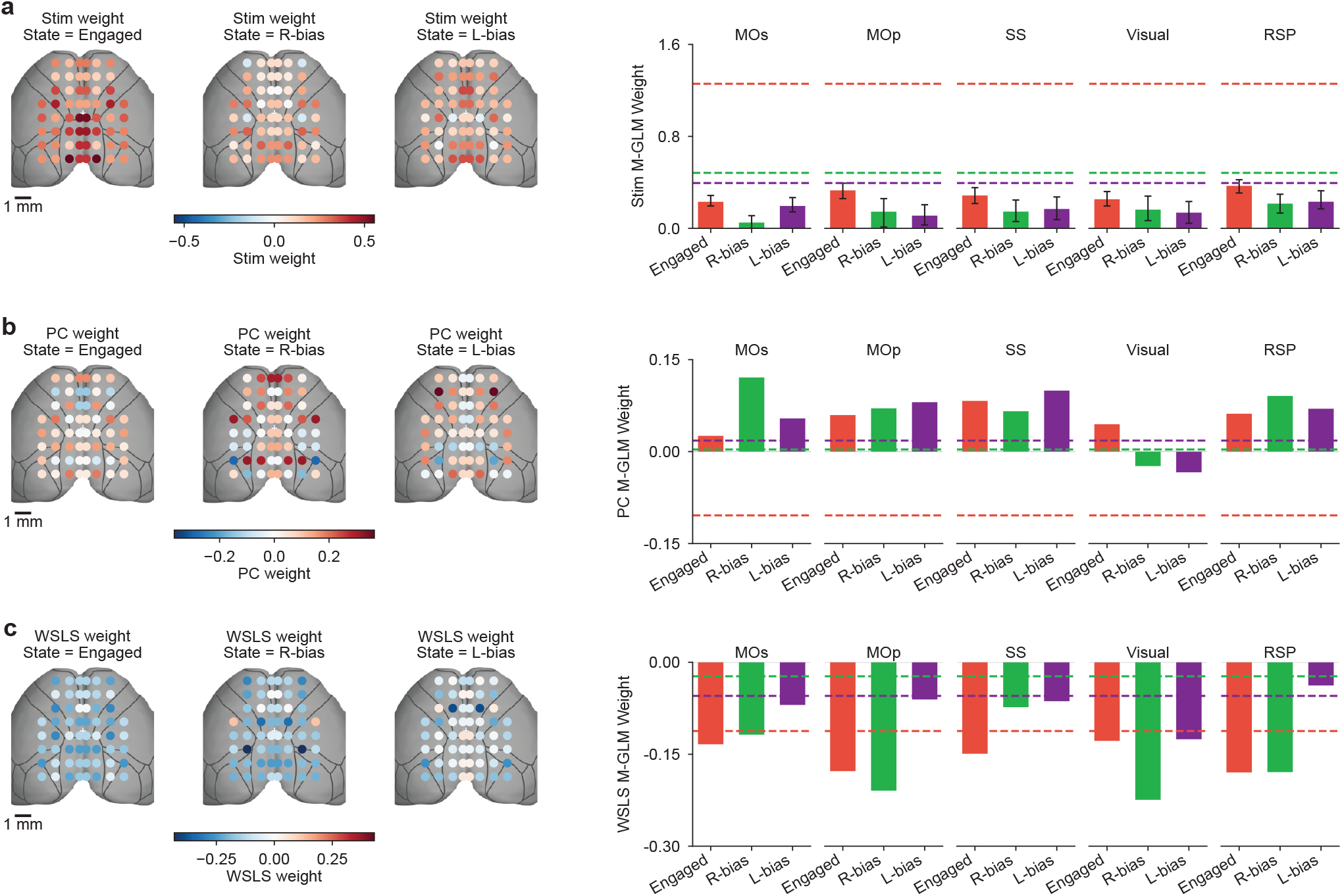
State-specific stimulus, previous-choice, and WSLS weights across cortical inactivation sites. **(a)** Cortical maps of stimulus weights (left) and summaries after grouping by anatomical region (right), separated by behavioral state. Weights are reduced relative to non-opto baselines (dashed lines) across regions. **(b)** Previous-choice (PC) weights. **(c)** Win-stay/lose-switch (WSLS) weights.

**Fig. S4:**
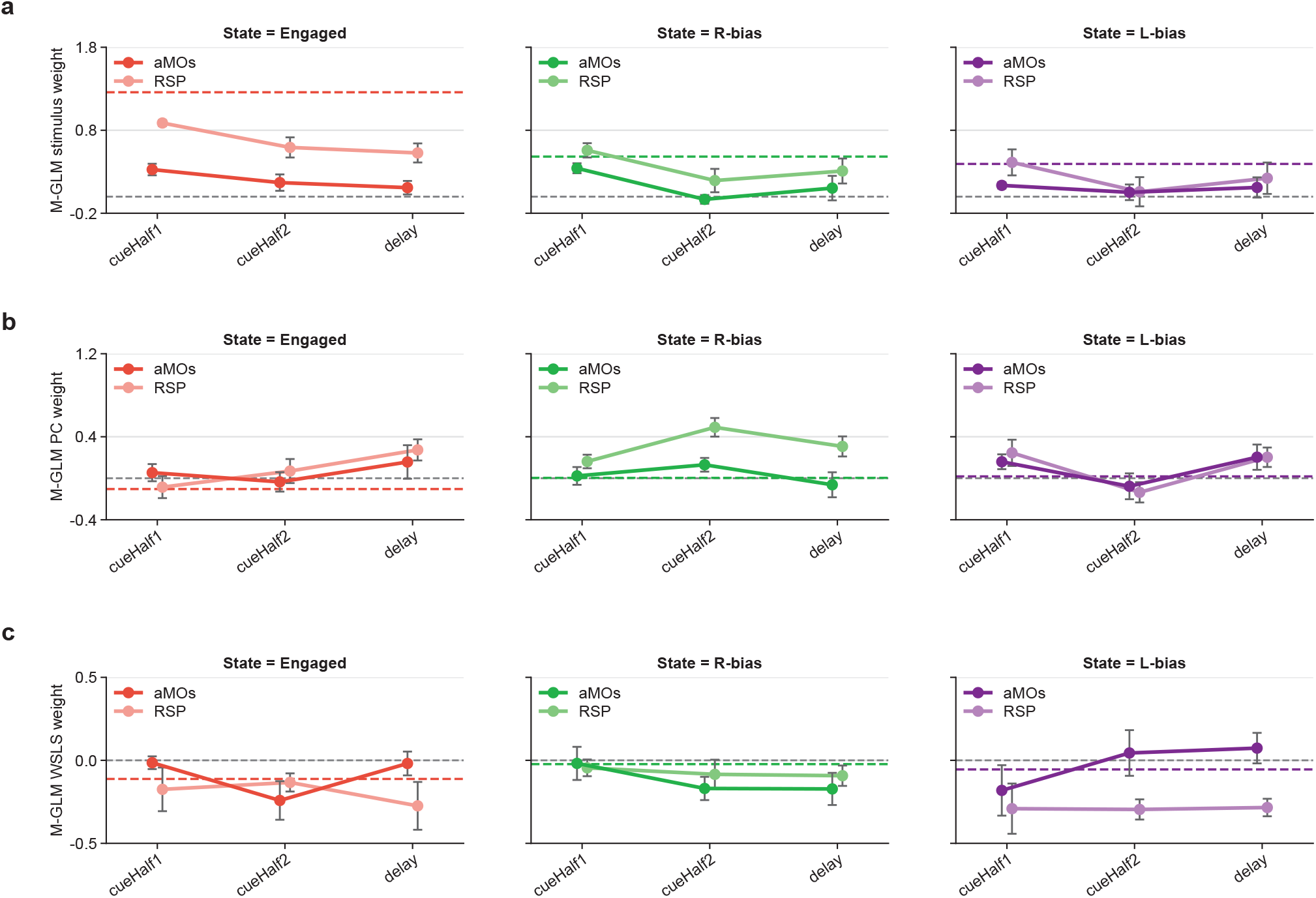
Subtrial cortical inactivation changes sensory and history-dependent choice weights across regions and epochs. **(a)** State-specific stimulus weights following aMOs or RSP inactivation during the first cue half, second cue half, or delay period. Stimulus weights tend to decrease across epochs (from cueHalf1 towards delay) and are generally larger following RSP than aMOs inactivation—results that are the inverse of the effects on bias seen in the main text. **(b)** Previous-choice (PC) weights. **(c)** Win-stay/lose-switch (WSLS) weights.

**Fig. S5:**
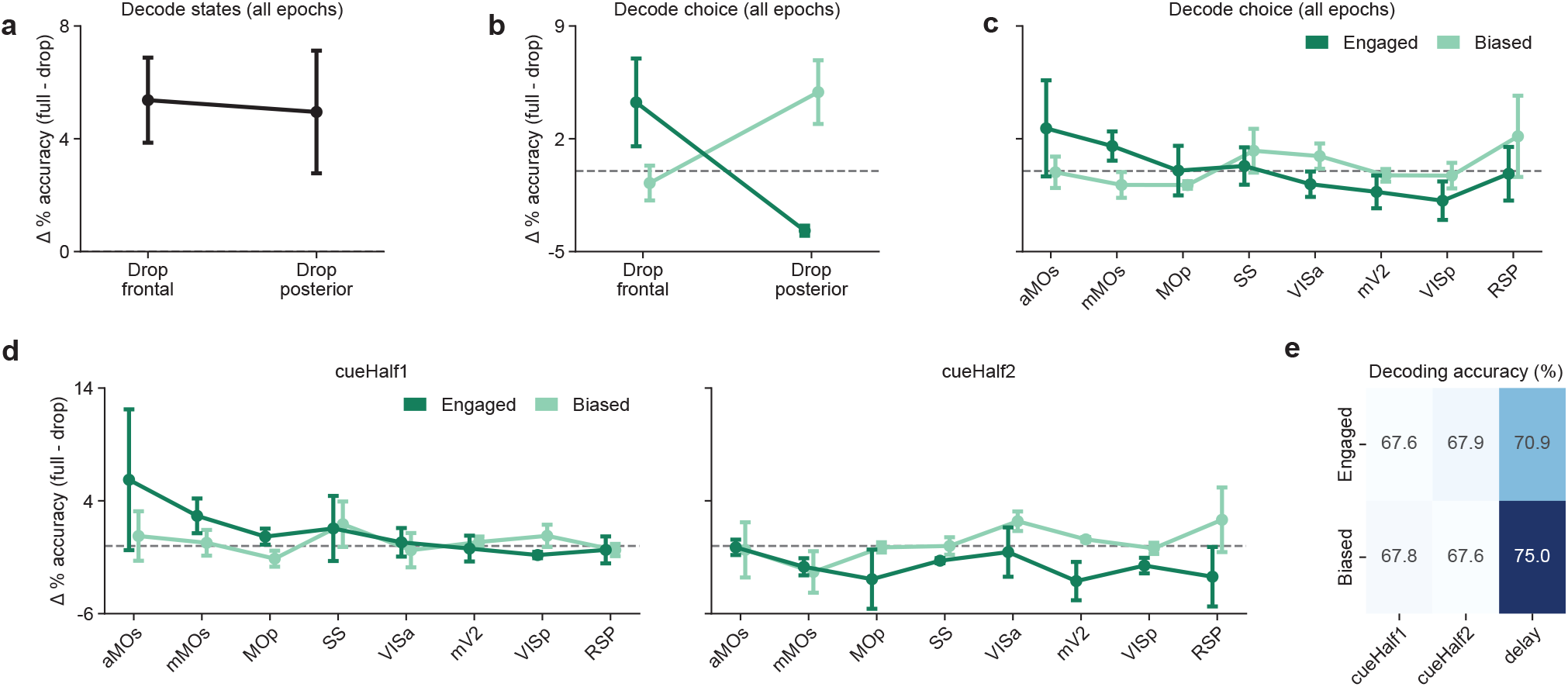
Regional contributions to state and choice decoding across maze epochs. **(a)** Decoding engaged versus biased states using activity averaged across all epochs: loss of decoding accuracy (full minus reduced model, percentage points; mean ± s.e.m. across mice) after excluding the frontal or posterior regions together. **(b)** Same as (a), for decoding choice (left versus right) within engaged and biased trials, using separate decoders. **(c)** As in (b), after excluding each cortical region on its own. **(d)** As in (c), using activity from the first (0–100 cm, left) and second (100–200 cm, right) cue halves only. The delay epoch (200–300 cm) is shown in Fig. 4. **(e)** Full-model choice-decoding accuracy for each state and epoch, showing that choice is most accurately decoded during the delay period in both states.

## References

1. Busse, L. et al. The Detection of Visual Contrast in the Behaving Mouse. Journal of Neuroscience 31, 11351–11361 (2011).

2. Pinto, L. & Dan, Y. Cell-Type-Specific Activity in Prefrontal Cortex during Goal-Directed Behavior. Neuron 87, 437–450 (2015).

3. Morcos, A. S. & Harvey, C. D. History-dependent variability in population dynamics during evidence accumulation in cortex. Nature Neuroscience 19, 1672–1681 (2016).

4. Scott, B. B. et al. Fronto-parietal cortical circuits encode accumulated evidence with a diversity of timescales. Neuron 95, 385–398.e5 (2017).

5. Hwang, E. J., Dahlen, J. E., Mukundan, M. & Komiyama, T. History-based action selection bias in posterior parietal cortex. Nature Communications 8, 1242 (2017).

6. Pinto, L. et al. An Accumulation-of-Evidence Task Using Visual Pulses for Mice Navigating in Virtual Reality. English. Frontiers in Behavioral Neuroscience 12. Publisher: Frontiers, 36. ISSN: 1662-5153. https://www.frontiersin.org/articles/10.3389/fnbeh.2018.00036/full?report=reader (2020) (2018).

7. Akrami, A., Kopec, C. D., Diamond, M. E. & Brody, C. D. Posterior parietal cortex represents sensory history and mediates its effects on behaviour. Nature 554, 368–372 (2018).

8. Pisupati, S., Chartarifsky-Lynn, L., Khanal, A. & Churchland, A. K. Lapses in perceptual decisions reflect exploration. eLife 10, e55490 (2021).

9. Bolkan, S. S. et al. Opponent control of behavior by dorsomedial striatal pathways depends on task demands and internal state. Nature Neuroscience 25, 345–357. 10.1038/s41593-022-01021-9 (2022).

10. Bandi, A. C. et al. Parietal cortex is causally required for state-dependent decisions. Cell Reports 44, 116672 (2025).

11. Roy, N. A. et al. Extracting the dynamics of behavior in sensory decision-making experiments. Neuron 109, 597–610.e6 (2021).

12. Ashwood, Z. C. et al. Mice alternate between discrete strategies during perceptual decisionmaking. Nature Neuroscience 25, 201–212 (2022).

13. Mohammadi, Z., Ashwood, Z. C., The International Brain Laboratory & Pillow, J. W. Identifying the factors governing internal state switches during nonstationary sensory decision-making. Nature Communications 16, 11684 (2025).

14. Yin, C. et al. Spontaneous movements and their relationship to neural activity fluctuate with latent engagement states. Neuron 113, 3048–3063.e5 (2025).

15. Luo, J. K. et al. A cholinergic mechanism orchestrating task-dependent computation across the cortex. bioRxiv. preprint, posted 2025-11-27; confirmed by Lucas Pinto as the correct paper for the cholinergic-mechanism sentence (2025).

16. Gilad, A., Gallero-Salas, Y., Groos, D. & Helmchen, F. Behavioral Strategy Determines Frontal or Posterior Location of Short-Term Memory in Neocortex. Neuron 99, 814–828.e7 (2018).

17. Oz Rokach, R., Marmor, O., Levy, Y. & Gilad, A. Brain-wide pathways for object identity and location are dominated by internal strategies. Nature Communications. volume/article number not yet listed in Crossref as of verification; confirm against published page (2026).

18. Pinto, L. et al. Task-dependent changes in the large-scale dynamics and necessity of cortical regions. Neuron 104, 810–824.e9 (2019).

19. Pinto, L., Tank, D. W. & Brody, C. D. Multiple timescales of sensory-evidence accumulation across the dorsal cortex. eLife 11, e70263 (2022).

20. Escola, S., Fontanini, A., Katz, D. & Paninski, L. Hidden Markov models for the stimulus-response relationships of multistate neural systems. Neural Computation 23, 1071–1132 (2011).

21. Odoemene, O., Pisupati, S., Nguyen, H. & Churchland, A. K. Visual Evidence Accumulation Guides Decision-Making in Unrestrained Mice. Journal of Neuroscience 38, 10143–10155 (2018).

22. Calhoun, A. J., Pillow, J. W. & Murthy, M. Unsupervised identification of the internal states that shape natural behavior. en. Nature Neuroscience 22. Number: 12 Publisher: Nature Publishing Group, 2040–2049. ISSN: 1546-1726. https://www.nature.com/articles/s41593-019-0533-x (2020) (Dec. 2019).

23. Beron, C. C., Neufeld, S. Q., Linderman, S. W. & Sabatini, B. L. Mice exhibit stochastic and efficient action switching during probabilistic decision making. Proceedings of the National Academy of Sciences 119, e2113961119 (2022).

24. Le, N. M. et al. Mixtures of strategies underlie rodent behavior during reversal learning. PLOS Computational Biology 19, e1011430 (2023).

25. Hulsey, D., Zumwalt, K., Mazzucato, L., McCormick, D. A. & Jaramillo, S. Decision-making dynamics are predicted by arousal and uninstructed movements. Cell Reports 43, 113709 (2024).

26. Hennig, J. A. et al. Learning is shaped by abrupt changes in neural engagement. Nature Neuroscience 24, 727–736 (2021).

27. Mohammadi, Z. et al. Identifying latent states in decision-making from cortical inactivation data. Computational and Systems Neuroscience (2022).

28. Li, B., Nguyen, T. P., Ma, C. & Dan, Y. Inhibition of impulsive action by projection-defined prefrontal pyramidal neurons. Proceedings of the National Academy of Sciences 117, 17278–17287 (2020).

29. Narayanan, N. S. & Laubach, M. Top-Down Control of Motor Cortex Ensembles by Dorsomedial Prefrontal Cortex. Neuron 52, 921–931 (2006).

30. Guo, Z. V. et al. Flow of cortical activity underlying a tactile decision in mice. Neuron 81, 179–194 (2014).

31. Goard, M. J., Pho, G. N., Woodson, J. & Sur, M. Distinct roles of visual, parietal, and frontal motor cortices in memory-guided sensorimotor decisions. eLife 5, e13764 (2016).

32. Barthas, F. & Kwan, A. C. Secondary Motor Cortex: Where ‘Sensory’ Meets ‘Motor’ in the Rodent Frontal Cortex. Trends in Neurosciences 40, 181–193 (2017).

33. Zatka-Haas, P., Steinmetz, N. A., Carandini, M. & Harris, K. D. Sensory coding and the causal impact of mouse cortex in a visual decision. eLife 10, e63163 (2021).

34. Arlt, C. et al. Cognitive experience alters cortical involvement in goal-directed navigation. eLife 11, e76051 (2022).

35. Gupta, D. et al. A multi-region recurrent circuit for evidence accumulation in rats. Neuron 114, 521–535.e5 (2026).

36. Steinmetz, N. A., Zatka-Haas, P., Carandini, M. & Harris, K. D. Distributed coding of choice, action and engagement across the mouse brain. Nature 576, 266–273 (2019).

37. Cazettes, F. et al. A reservoir of foraging decision variables in the mouse brain. Nature Neuroscience 26, 840–849 (2023).

38. Khilkevich, A. et al. Brain-wide dynamics linking sensation to action during decision-making. Nature 634, 890–900 (2024).

39. Bondy, A. et al. Multi-region coordination of evidence accumulation and decision commitment in the brain. bioRxiv. preprint, posted 2026-07-09 – not yet confirmed as journal-accepted (2026).

40. Lebedeva, A. et al. Dorsal prefrontal cortex drives perseverative behavior in mice. Nature Communications 17, 5715 (2026).

41. Hattori, R., Danskin, B., Babic, Z., Mlynaryk, N. & Komiyama, T. Area-Specificity and Plasticity of History-Dependent Value Coding During Learning. Cell 177, 1858–1872.e15 (2019).

42. Zhang, Y. E. et al. Adaptive reorganization of history encoding in the retrosplenial cortex supports flexible decision-making strategies. Neuron 114, 2272–2285.e6 (2026).

43. Wallis, J. D., Anderson, K. C. & Miller, E. K. Single neurons in prefrontal cortex encode abstract rules. Nature 411, 953–956 (2001).

44. Ridley, R. M., Durnford, L. J., Baker, J. A. & Baker, H. F. Cognitive inflexibility after archicortical and paleocortical prefrontal lesions in marmosets. Brain Research 628, 56–64 (1993).

45. Rodgers, C. C. & DeWeese, M. R. Neural correlates of task switching in prefrontal cortex and primary auditory cortex in a novel stimulus selection task for rodents. Neuron 82, 1157–1170 (2014).

46. Cole, N., Harvey, M., Myers-Joseph, D., Gilra, A. & Khan, A. G. Prediction-error signals in anterior cingulate cortex drive task-switching. Nature Communications 15, 7088 (2024).

47. Bouret, S. & Sara, S. J. Network reset: a simplified overarching theory of locus coeruleus noradrenaline function. Trends in Neurosciences 28, 574–582 (2005).

48. Prado, V. F., Janickova, H., Al-Onaizi, M. A. & Prado, M. A. M. Cholinergic circuits in cognitive flexibility. Neuroscience 345, 130–141 (2017).

49. Tervo, D. G. R. et al. Behavioral Variability through Stochastic Choice and Its Gating by Anterior Cingulate Cortex. Cell 159, 21–32 (2014).

50. Wang, H. et al. Frontal noradrenergic and cholinergic transients exhibit distinct spatiotemporal dynamics during competitive decision-making. Science Advances 11, eadr9916 (2025).

51. McBurney-Lin, J., Vargova, G., Garad, M., Zagha, E. & Yang, H. The locus coeruleus mediates behavioral flexibility. Cell Reports 41, 111534 (2022).

52. Berman, G. J., Choi, D. M., Bialek, W. & Shaevitz, J. W. Mapping the stereotyped behaviour of freely moving fruit flies. Journal of The Royal Society Interface 11, 20140672 (2014).

53. Wiltschko, A. B. et al. Mapping sub-second structure in mouse behavior. Neuron 88, 1121–1135 (2015).

54. Eyjolfsdottir, E., Branson, K., Yue, Y. & Perona, P. Learning recurrent representations for hierar-chical behavior modeling. International Conference on Learning Representations (ICLR) (2017).

55. Katsov, A. Y., Freifeld, L., Horowitz, M., Kuehn, S. & Clandinin, T. R. Dynamic structure of locomotor behavior in walking fruit flies. Elife 6, e26410 (2017).

56. Mohammadi, Z. et al. Identifying latent states in decision-making from cortical inactivation data. Computational and Systems Neuroscience (2022).

57. Li, J.-J., Shi, C., Li, L. & Collins, A. G. Dynamic noise estimation: A generalized method for modeling noise fluctuations in decision-making. Journal of Mathematical Psychology 119, 102842 (2024).

58. Bishop, C. M. Pattern recognition and machine learning (springer, 2006).

59. Dempster, A. P., Laird, N. M. & Rubin, D. B. Maximum likelihood from incomplete data via the EM algorithm. Journal of the royal statistical society: series B (methodological) 39, 1–22 (1977).

60. Bengio, Y. & Frasconi, P. An input output HMM architecture. Advances in neural information processing systems 7 (1994).

61. Linderman, S., Antin, B., Zoltowski, D. & Glaser, J. SSM: Bayesian learning and inference for state space models https://github.com/lindermanlab/ssm. software package; specific version used not independently confirmed. 2020.

62. Salakhutdinov, R., Roweis, S. T. & Ghahramani, Z. Optimization with EM and expectation-conjugategradient in Proceedings of the 20th International Conference on Machine Learning (ICML-03) (2003), 672–679.

63. Baum, L. E., Petrie, T., Soules, G. & Weiss, N. A maximization technique occurring in the statistical analysis of probabilistic functions of Markov chains. The annals of mathematical statistics 41, 164– 171 (1970).

64. Virtanen, P. et al. SciPy 1.0: Fundamental Algorithms for Scientific Computing in Python. Nature Methods 17, 261–272 (2020).

